# Closely related type II-C Cas9 orthologs recognize diverse PAMs

**DOI:** 10.1101/2022.02.21.481263

**Authors:** Jingjing Wei, Linghui Hou, Jingtong Liu, Siqi Gao, Tao Qi, Shuna Sun, Yongming Wang

## Abstract

Autosomal dominant diseases can be treated by allele-specific disruption of mutant alleles. If the missense mutation generates a novel protospacer-adjacent motif (PAM), CRISPR-Cas9 can distinguish mutant alleles from wild-type alleles by a PAM-specific approach. Therefore, it is crucial to develop a CRISPR toolbox capable of recognizing multiple PAMs. Here, by using a GFP-activation assay, we tested the activities of 29 type II-C orthologs closely related to Nme1Cas9, 25 of which are active in human cells. These orthologs recognize diverse PAMs with variable length and nucleotide preference, including purine-rich, pyrimidine-rich, and mixed purine and pyrimidine PAMs. We characterized in depth the activity and specificity of Nsp2Cas9. We also generated a chimeric Cas9 nuclease that recognizes a simple N_4_C PAM, representing the most relaxed PAM preference for compact Cas9s to date. These Cas9 nucleases significantly enhance our ability to perform allele-specific genome editing.

## Introduction

RNA-guided CRISPR/Cas9 was originally identified as part of the microbial adaptive immune system, which contains three crucial components: a Cas9 endonuclease, a CRISPR RNA (crRNA) and a trans-activating CRISPR RNA (tracrRNA) (1, 2). These three components constitute an active ribonucleoprotein complex to recognize and cleave foreign DNA (3, 4). To recognize foreign DNA, the target sequence should contain (i) a complementary sequence with the crRNA and (ii) a protospacer-adjacent motif (PAM) immediately downstream of the target (4). The PAM allows this system to differentiate between the DNA target in invading genetic material (non-self) and the same DNA sequence encoded within CRISPR arrays (self) (5).

The competitive coevolution of CRISPR/Cas9 systems with different evolving viruses leads to acceptance of diverse PAMs across the Cas9 nucleases (6, 7). For instance, SpCas9 recognizes an NGG PAM (8-11); SaCas9 recognizes an NNGRRT PAM (R=A, G) (12); CjCas9 recognizes an N4RYAC (Y=C, T) PAM (13); St1Cas9 recognizes an NNRGAA PAM (14); BlatCas9 recognizes an N_4_CNAA PAM (15). A recent study screened a list of 79 Cas9 orthologs and identified multiple PAMs, including A-, C-, T-, and G-rich nucleotides, from single nucleotide recognition to sequence strings longer than 4 nucleotides (7). Nevertheless, these orthologs are phylogenetically distinct and whether they are active in human cells remains unknown.

The existing evidence reveals that even phylogenetically closely related Cas9 orthologs may recognize distinct PAMs. In this study, “closely related Cas9 orthologs” means that these orthologs have >50% sequence identity and can recognize each other’s processed crRNA:tracrRNA duplexes as the guide RNA (gRNA). SpCas9 and SaCas9 are representative type II-A Cas9 nucleases (16-18). ScCas9 has 83.3% sequence identity with SpCas9 and recognizes an NNG PAM (17). SmacCas9 has 58.4% sequence identity with SpCas9 and recognizes an NAA PAM (19). We recently identified four Cas9 orthologs that are closely related to SaCas9 and recognized NNGG, NNGRR, and NNGGV PAMs (V=A, C, G), respectively (20, 21). However, type II-C Cas9 nucleases account for nearly half of the total type II Cas9s (22), but their PAM diversity within closely related orthologs remains largely unknown.

Several studies have demonstrated that CRISPR/Cas9 can treat autosomal dominant diseases through allele-specific disruption of mutant alleles. Autosomal dominant diseases are mainly caused by single-nucleotide missense mutations (23). If missense mutations form novel PAMs, CRISPR/Cas9 nucleases can disrupt mutant alleles by a PAM-specific approach (24, 25). Therefore, it is crucial to develop a CRISPR toolbox capable of recognizing multiple PAMs.

Nme1Cas9 is a representative type II-C ortholog that was first developed for genome editing in 2013 (26, 27). It is a compact and high-fidelity enzyme but recognizes a long PAM. To expand the targeting scope, Edraki et al. investigated two Nme1Cas9 orthologs (Nme2Cas9 and Nme3Cas9) and demonstrated that Nme2Cas9 recognized a simple N_4_CC PAM (18). In this study, we investigated the editing capacity of 29 Nme1Cas9 orthologs, 25 of which were active in human cells. These orthologs recognized diverse PAMs with variable length and nucleotide preference. Importantly, based on these orthologs, we generated a chimeric Cas9 nuclease that recognized a simple N_4_C PAM for genome editing. These Cas9 nucleases significantly enhance our ability to perform allele-specific genome editing.

## Results

### Investigation of PAM diversity within Nme1Cas9 orthologs

To investigate the PAM diversity within Nme1Cas9 orthologs, we selected 29 Nme1Cas9 orthologs from the UniProt database (28) using Nme1Cas9 as a reference (Figure 1-figure supplement 1 and Table 1). The amino-acid identities to Nme1Cas9 varied from 59.6% to 70.4%. A previous structural study had shown that residues Q981, H1024, T1027 and N1029 in the Nme1Cas9 PAM-interacting (PI) domain are crucial for PAM recognition (29). Amino acid sequence alignment revealed that 28 selected orthologs differed in at least one residue corresponding to the four residues of Nme1Cas9 (Figure 1-figure supplement 2A-C), indicating that these orthologs may recognize distinct PAMs. BdeCas9 had the same four residues as Nme1Cas9. Nme1Cas9 H1024 forms hydrogen bonds with the fifth nucleotide of the N_4_GATT PAM (29). According to the amino acids corresponding to the Nme1Cas9 H1024, the orthologs selected here could be divided into three groups, which contained aspartate (D), histidine (H), and asparagine (N) residues, respectively (Figure 1-figure supplement 1 2A-C).

**Table 1.** Nme1Cas9 orthologs selected from UniProt database.

| UniProt ID | Host strain | Name | Length (aa) | Identity to Nme1Cas9 (%) |
| --- | --- | --- | --- | --- |
| A0A011P7F8 | *Mannheimia granulomatis* | MgrCas9 | 1049 | 65.5 |
| A0A0A2YBT2 | *Gallibacterium anatis* IPDH697-78 | GanCas9 | 1035 | 59.7 |
| A0A0J0YQ19 | *Neisseria arctica* | NarCas9 | 1070 | 70.4 |
| A0A1T0B6J6 | *[Haemophilus felis* | HfeCas9 | 1058 | 65.3 |
| A0A1X3DFB7 | *Neisseria dentiae* | NdeCas9 | 1074 | 66.4 |
| A0A263HCH5 | *Actinobacillus seminis* | AseCas9 | 1059 | 66 |
| A0A2M8S290 | *Conservatibacter flavescens* | CflCas9 | 1063 | 64.2 |
| A0A2U0SK41 | *Pasteurella langaaensis* DSM 22999 | PlaCas9 | 1056 | 63.9 |
| A0A356E7S3 | *Pasteurellaceae bacterium* | PstCas9 | 1076 | 63 |
| A0A369Z1C7 | *Haemophilus parainfluenzae* | Hpa1Cas9 | 1056 | 64.8 |
| A0A369Z3K3 | *Haemophilus parainfluenzae* | Hpa2Cas9 | 1054 | 65.2 |
| A0A377J007 | *Haemophilus pittmaniae* | HpiCas9 | 1053 | 65.2 |
| A0A378UFN0 | *Bergeriella denitrificans* (*Neisseria denitrificans*) | BdeCas9 | 1069 | 68.8 |
| A0A379B6M0 | *Pasteurella mairii* | PmaCas9 | 1061 | 63.1 |
| A0A379CB86 | *Phocoenobacter uteri* | PutCas9 | 1059 | 63 |
| A0A380MYP0 | *Suttonella indologenes* | SinCas9 | 1071 | 67.8 |
| A0A3N3EE71 | *Neisseria animalis* | Nan1Cas9 | 1074 | 66.6 |
| A0A3S4XT82 | *Neisseria animaloris* | Nan2Cas9 | 1078 | 65.4 |
| A0A420XER8 | *Otariodibacter oris* | OorCas9 | 1058 | 64.4 |
| A0A448K7T0 | *Pasteurella aerogenes* | PaeCas9 | 1056 | 68.2 |
| A0A4S2QB06 | *Rodentibacter pneumotropicus* | RpnCas9 | 1055 | 63.7 |
| A0A4Y9GBC9 | *Neisseria sp. WF04* | Nsp2Cas9 | 1067 | 59.6 |
| A6VLA7 | *Actinobacillus succinogenes*<br>(strain ATCC 55618/DSM<br>22257/130Z) | AsuCas9 | 1062 | 64.6 |
| C5S1N0 | *Actinobacillus minor* NM305 | AmiCas9 | 1056 | 67.7 |
| E0F2V7 | *Actinobacillus pleuropneumoniae*<br>serovar 10 str. D13039 | ApsCas9 | 1054 | 65.4 |
| F2B8K0 | *Neisseria bacilliformis* ATCC BAA-1200 | NbaCas9 | 1077 | 66.5 |
| J4KDT3 | *Haemophilus sputorum* | HspCas9 | 1052 | 65.2 |
| V9H606 | *Simonsiella muelleri* ATCC 29453 | SmuCas9 | 1063 | 62.2 |
| W0Q6X6 | *Mannheimia* sp. USDA-ARS-USMARC-1261 | MspCas9 | 1047 | 65.7 |

Next, we investigated the conservation of the CRISPR direct repeat sequences and tracrRNA sequences among Nme1Cas9 orthologs. Both direct repeats and putative tracrRNAs were identified for 26 Cas9 orthologs. Sequence alignment revealed that direct repeats and the 5’ end of tracrRNAs were conserved among Nme1Cas9 orthologs (Figure 1-figure supplement 3A-B). We generated single guide RNA (sgRNA) scaffolds for these orthologs by fusing the 3′ end of a truncated direct repeat with the 5′ end of the corresponding tracrRNA, including the full-length tail, via a 4-nt linker. The RNAfold web server predicted that these sgRNAs contained a conserved stem loop at each end, similar to the Nme1Cas9 sgRNA (Figure 1-figure supplement 4). Twenty orthologs contained one stem loop in the middle, while six orthologs contained two stem loops in the middle.

Next, the human-codon-optimized Nme1Cas9 orthologs were synthesized and cloned into the Nme2Cas9_AAV plasmid backbone (18). We used a previously developed PAM-screening assay (20) to determine their PAMs. This is a GFP-activation assay where a 24-nt protospacer followed by an 8-bp random sequence is inserted between the ATG start codon and GFP-coding sequence, resulting in a frameshift mutation. The library was stably integrated into the human genome (HEK293T cells) using a lentivirus. If a Cas9 ortholog enables editing the protospacer, it will generate insertions/deletions (indels) at the protospacer and induce GFP expression in a portion of cells (Figure 1A-B). The Nme1Cas9 sgRNA scaffold was used for all Cas9 orthologs in this study. Each Cas9 expression plasmid was co-transfected with an sgRNA plasmid into the cell library. The cells without transfection were included as a negative control, and Nme2Cas9 was included as a positive control. Five days after transfection, GFP-positive cells were observed for 25 orthologs through fluorescence microscope. The percentage of GFP-positive cells varied from ∼0.01% to 0.18%, as revealed by flow cytometry analysis (Figure 1C).

**Figure 1.**
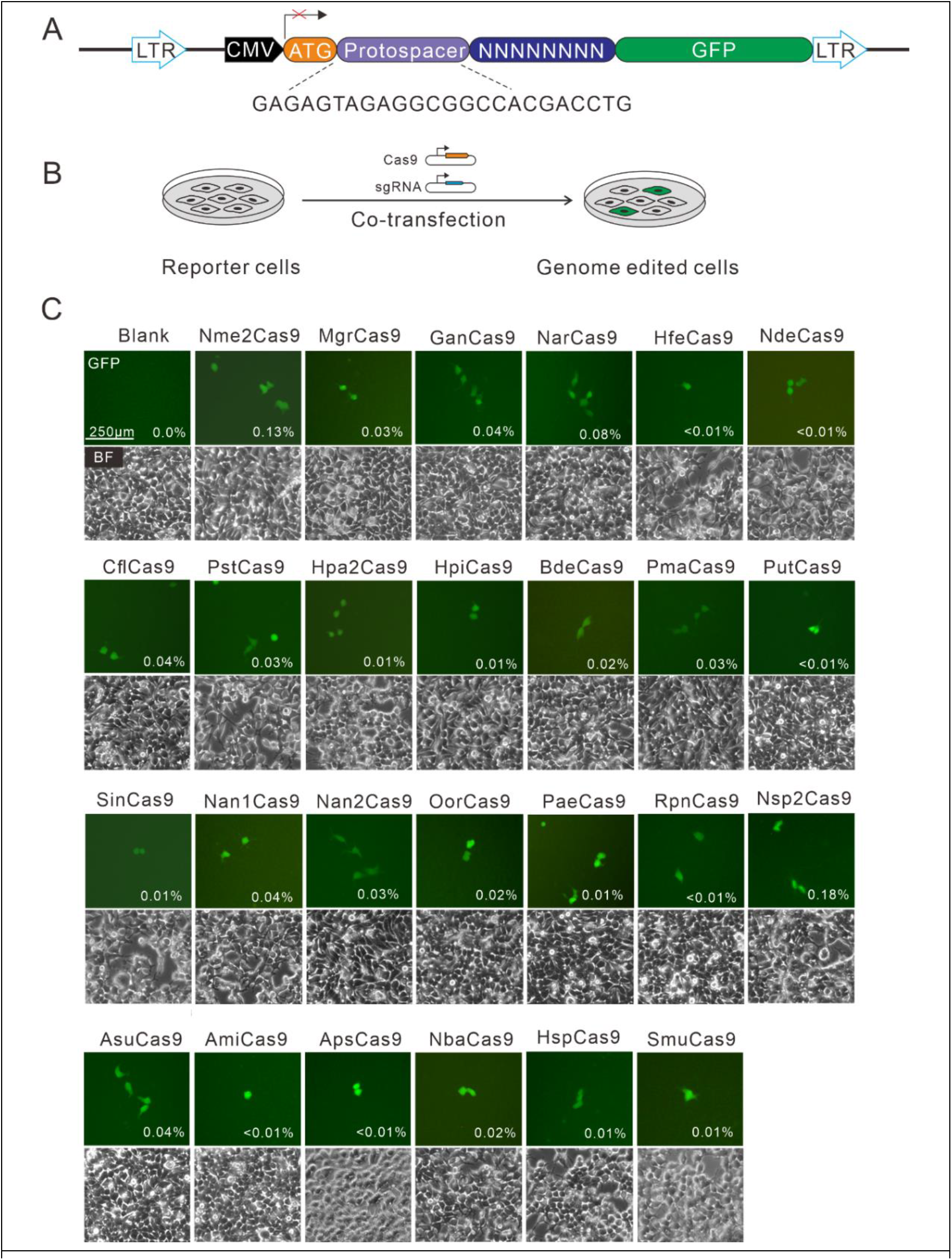
Screening of Nme1Cas9 orthologs activities through a GFP-activation assay. **(A)** Schematic of the GFP-activation assay. A protospacer flanked by an 8-bp random sequence is inserted between the ATG start codon and GFP-coding sequence, resulting in a frameshift mutation. The library DNA is stably integrated into HEK293T cells via lentivirus infection. Genome editing can lead to in-frame mutation. The protospacer sequence is shown below. **(B)** Procedure of the GFP-activation assay. Cas9 and sgRNA expression plasmids were co-transfected into the reporter cells. GFP-positive cells could be observed if the protospacer is edited. **(C)** 25 out of 29 Nme1Cas9 orthologs could induce GFP expression. The percentage of GFP-positive cells is shown. Reporter cells without Cas9 transfection are used as a negative control. Scale bar: 250 μm.

Next, we analyzed the PAM for each active ortholog. The GFP-positive cells were sorted by flow cytometry, and the protospacer and the 8-bp random sequence were PCR-amplified for deep sequencing. The sequencing results revealed that indels were generated at the target site (Figure 2A). We generated the WebLogo diagram for each ortholog based on deep sequencing data. Typically, Nme1Cas9 orthologs displayed minimal or no base preference at PAM positions 1-4 (Figure 2-figure supplement 1A-B). For the orthologs that potentially require PAMs longer than 8-bp, we shifted the target sequence by three nucleotides in the 5’ direction to allow PAM identification to be extended from 8 to 11 bp.

**Figure 2.**
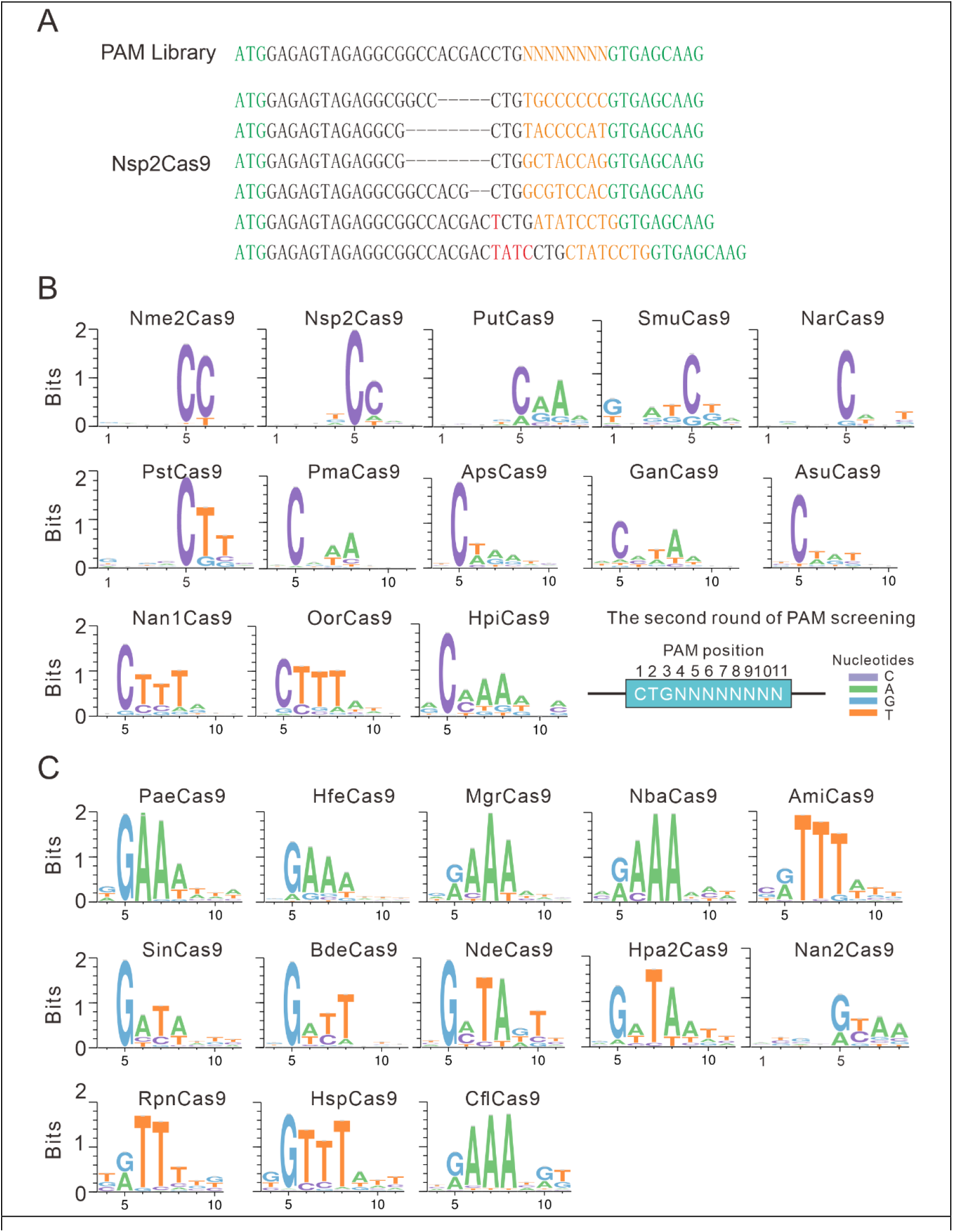
PAM analysis for each Cas9 nuclease. **(A)** Example of indel sequences measured by deep sequencing for Nsp2Cas9. The GFP coding sequences are shown in green; an 8-bp random sequence is shown in orange; black dashes indicate deleted bases; red bases indicate insertion mutations. **(B)** The PAM WebLogos for Nme1Cas9 orthologs containing an aspartate residue corresponding to the Nme1Cas9 H1024. PAM positions for each WebLogo are shown below. The PAM WebLogos for Nme2Cas9, Nsp2Cas9, PutCas9, SmuCas9, NarCas9, PstCas9 are generated from the first round of PAM screening and the PAM WebLogos for others are generated from the second round of PAM screening. PAM positions for the second round of screening are shown on the bottom right. (**C**) The PAM WebLogos for Nme1Cas9 orthologs containing histidine, or asparagine residues corresponding to the Nme1Cas9 H1024. PAM positions for each WebLogo are shown below. The PAM WebLogo for Nan2Cas9 is generated from the first round of PAM screening and the PAM WebLogos for others are generated from the second round of PAM screening.

Interestingly, Nme1Cas9 orthologs recognized diverse PAMs (Figure 2B-C and Figure 2-figure supplement 2). Generally, they displayed strong nucleotide preferences for up to four base pairs. When orthologs contained an aspartate (D) residue corresponding to the Nme1Cas9 H1024, they displayed a strong C preference at PAM position 5 (Figure 2B); when orthologs contained histidine (H), or asparagine (N) residues corresponding to the Nme1Cas9 H1024, they displayed a strong R (R=A or G) preference at PAM position 5 (Figure 2C). The length of PAM recognition varied between 5 and 11 base pairs. Nme2Cas9 recognized an N_4_CC PAM, consistent with a previous report (18). BdeCas9 recognized an N_4_GATT PAM that was the same as Nme1Cas9 (26). In addition, many orthologs displayed degenerate PAM recognition (e.g., HpiCas9, MgrCas9, and NbaCas9). Since Nsp2Cas9 induced more GFP-positive cells (0.18%) than other Cas9s and recognized a simple N_4_CC PAM, we focused on Nsp2Cas9 in the following study.

### Nsp2Cas9 enables genome editing for endogenous sites

To test whether Nsp2Cas9 enables genome editing for endogenous targets in human cells, we selected a panel of 19 targets. Nme2Cas9 has the same expression backbone as Nsp2Cas9 and was used for side-by-side comparison (Figure 3A). Western blot analysis revealed that their protein expression levels were comparable (Figure 3B). Five days after transfection of Cas9 and sgRNA expression plasmids, cells were harvested, and genomic DNA was extracted for targeted deep sequencing. The results revealed that Nsp2Cas9 and Nme2Cas9 displayed comparable editing activity, although the activities varied depending on the targets (Figure 3C-D). We tested Nsp2Cas9 editing capacity in additional cell lines, including HeLa, HCT116, A375, SH-SY5Y, and mouse N2a cells, and it could also generate indels in these cells with varying efficiencies (Figure 3-figure supplement 1A-E). In summary, these results demonstrated that Nsp2Cas9 could serve as a new player for genome editing.

**Figure 3.**
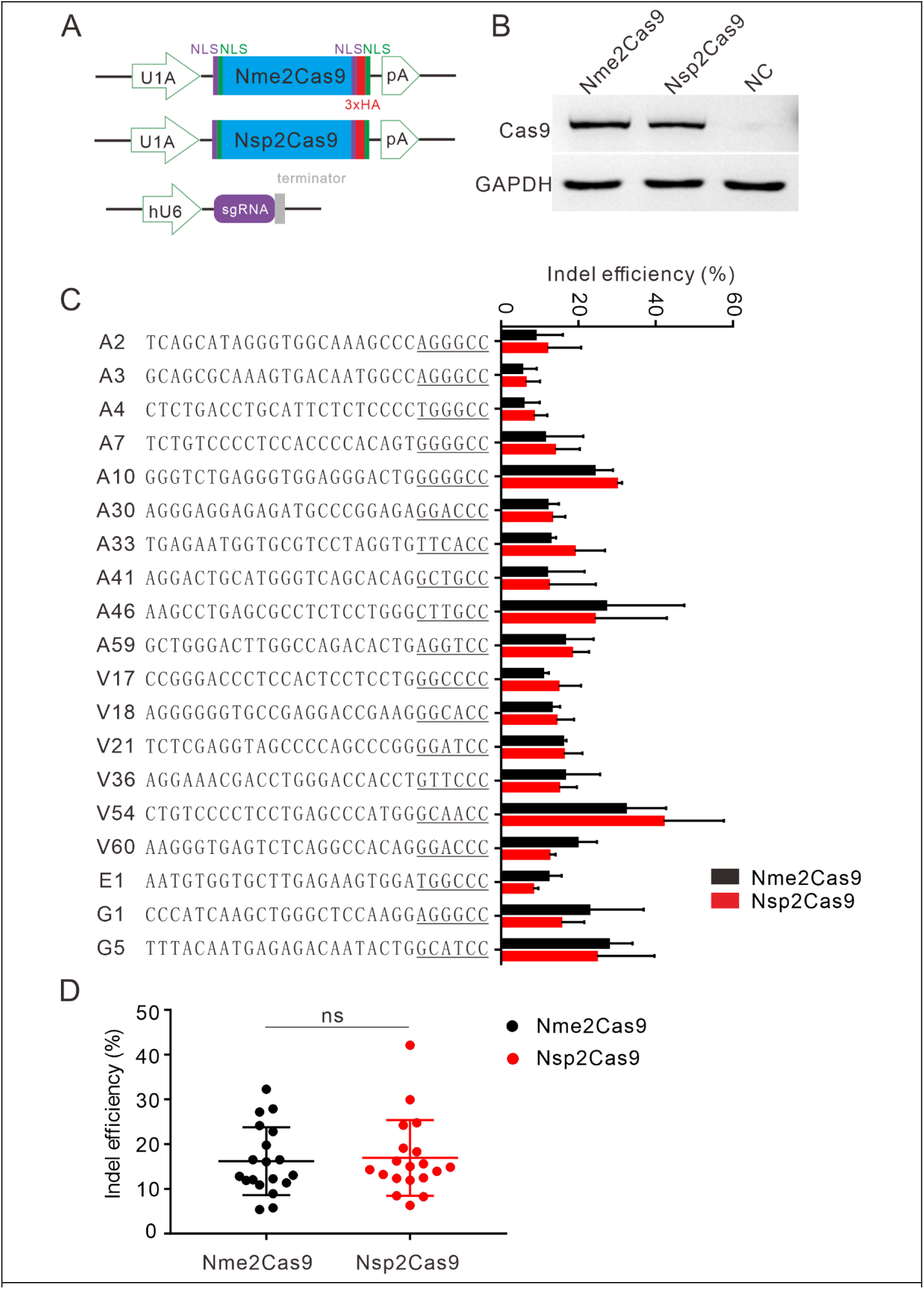
Nsp2Cas9 enables editing in HEK293T cells. **(A)** Schematic of Cas9 and sgRNA expression constructs. U1A: U1A promoter; pA: polyA; NLS: nuclear localization signal; HA: HA tag. **(B)** Protein expression levels of Nsp2Cas9 and Nme2Cas9 were analyzed by Western blot. HEK293T cells without Cas9 transfection were used as negative control (NC). **(C)** Comparison of Nsp2Cas9 and Nme2Cas9 editing efficiencies at 19 endogenous loci in HEK293T cells. Data represent mean ± SD for n=3 biologically independent experiments. **(D)** Quantification of the indel efficiencies for Nsp2Cas9 and Nme2Cas9. Each dot represents an average efficiency for an individual locus. Data represent mean ± SD. *P* values were determined using a two-sided Student’s t test. *P* =0.7486 (*P* >0.05), ns stands for not significant.

It has been reported that residues S593 and W596 in Nme1Cas9 play vital roles in cleavage activity. Replacement of the single or double residues with arginine could increase the cleavage activity of Nme1Cas9 in vitro (29). To test whether these two residues could influence Nsp2Cas9 activity, we identified the corresponding residues S597 and W600 in Nsp2Cas9 by protein sequence alignment and replaced the single or double residues with arginine. To test the activities of the resulting Cas9 variants, we constructed a GFP-activation cell line that was similar to the PAM-screening construct but with a fixed PAM (Figure 3-figure supplement 2A). The editing activities could be reflected by the GFP-positive cells. However, five days after genome editing, the results showed that single or double mutation variants could not improve the editing activities (Figure 3-figure supplement 2B).

### NarCas9 enables genome editing for endogenous sites

In addition to Nsp2Cas9, we also tested the editing ability of NarCas9, which recognizes a simple N_4_C PAM. Five days after transfection of NarCas9 and sgRNA expression plasmids, the cells were harvested, and genomic DNA was extracted for targeted deep sequencing. The results showed that NarCas9 could generate indels in both HEK293T and HeLa cells, but the efficiency was low (Figure 3-figure supplement 3A-C). SmuCas9 also recognizes a simple N_4_C PAM, but it exhibited minimal activity in the PAM-screening assay, and we did not study this further.

### A chimeric Cas9 nuclease enables genome editing for endogenous sites

We and others have demonstrated that swapping the PI domain between closely related orthologs can generate a chimeric nuclease that recognizes distinct PAMs (18-20). To expand the targeting scope, we replaced the Nsp2Cas9 PI with SmuCas9 PI, resulting in a chimeric Cas9 nuclease that we named “Nsp2-SmuCas9’’ (Figure 4A). The PAM-screening assay revealed that only Nsp2-SmuCas9 could induce GFP expression. Deep sequencing analysis revealed that Nsp2-SmuCas9 recognized an N_4_C PAM (Figure 4B-C). We tested the activities of Nsp2-SmuCas9 in a panel of 12 endogenous targets that have been used for NarCas9. Targeted deep sequencing revealed that Nsp2-SmuCas9 could efficiently induce indels at these sites (Figure 4D). Importantly, Nsp2-SmuCas9 displayed higher activities than NarCas9 (Figure 4E). We also replaced the Nsp2Cas9 PI with NarCas9 PI, resulting in a chimeric Cas9 nuclease that we named “Nsp2-NarCas9’’. However, Nsp2-NarCas9 showed no activity in the PAM-screening assay.

**Figure 4.**
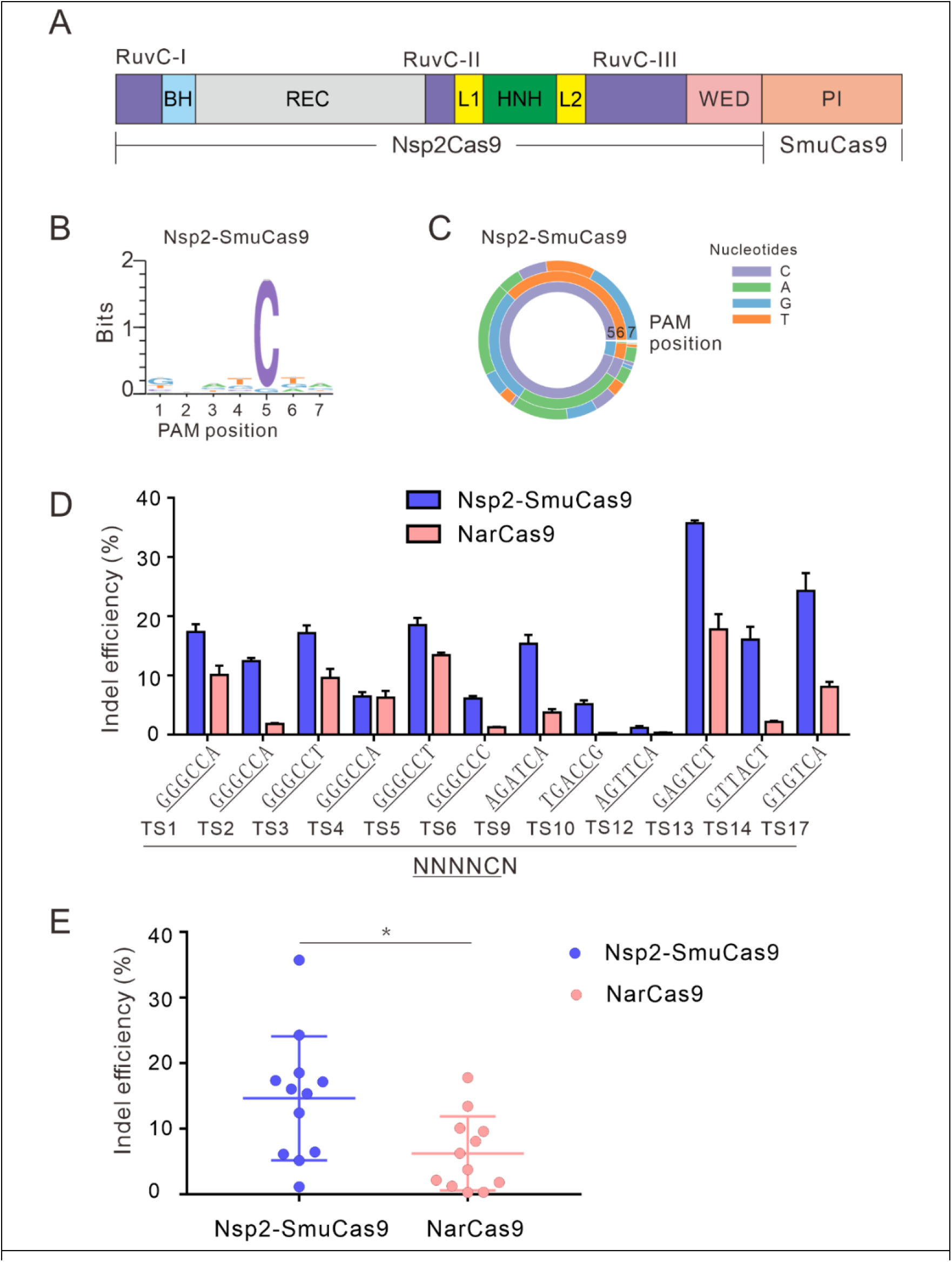
Characterization of Nsp2-SmuCas9 for genome editing. **(A)** Schematic diagram of chimeric Cas9 nucleases based on Nsp2Cas9. PI domain of Nsp2Cas9 was replaced with the PI domain of SmuCas9. **(B)** Sequence logos and **(C)** PAM wheel diagrams indicate that Nsp2-SmuCas9 recognizes an N_4_C PAM. **(D)** Nsp2-SmuCas9 generated indels at endogenous sites with N_4_C PAMs in HEK293T cells. Indel efficiencies were determined by targeted deep sequencing. NarCas9 is used as a control. Data represent mean ± SD for n=3 biologically independent experiments. **(E)** Quantification of the indel efficiencies for Nsp2-SmuCas9 and NarCas9. Each dot represents an average efficiency for an individual locus. Data represent mean ± SD. *P* values were determined using a two-sided Student’s *t* test. \**P* =0.0148 (0.01< *P* <0.05).

### Analysis of Nsp2Cas9 and Nsp2-SmuCas9 specificity

Next, we analyzed Nsp2Cas9 specificity by using the GFP-activation assay, and Nme2Cas9 was used for comparison. We generated a panel of 11 sgRNAs with dinucleotide mismatches. Five days after co-transfection of Nsp2Cas9 with individual sgRNAs, GFP-positive cells were analyzed by using a flow cytometer. The results showed that Nme2Cas9 was a highly specific enzyme that did not tolerate dinucleotide mismatches (sgRNAs M2-M11), consistent with a previous study (18). In contrast, Nsp2Cas9 displayed moderate off-target effects with some sgRNAs (sgRNAs M1, M4, M8, M9, and M10) (Figure 5A).

**Figure 5.**
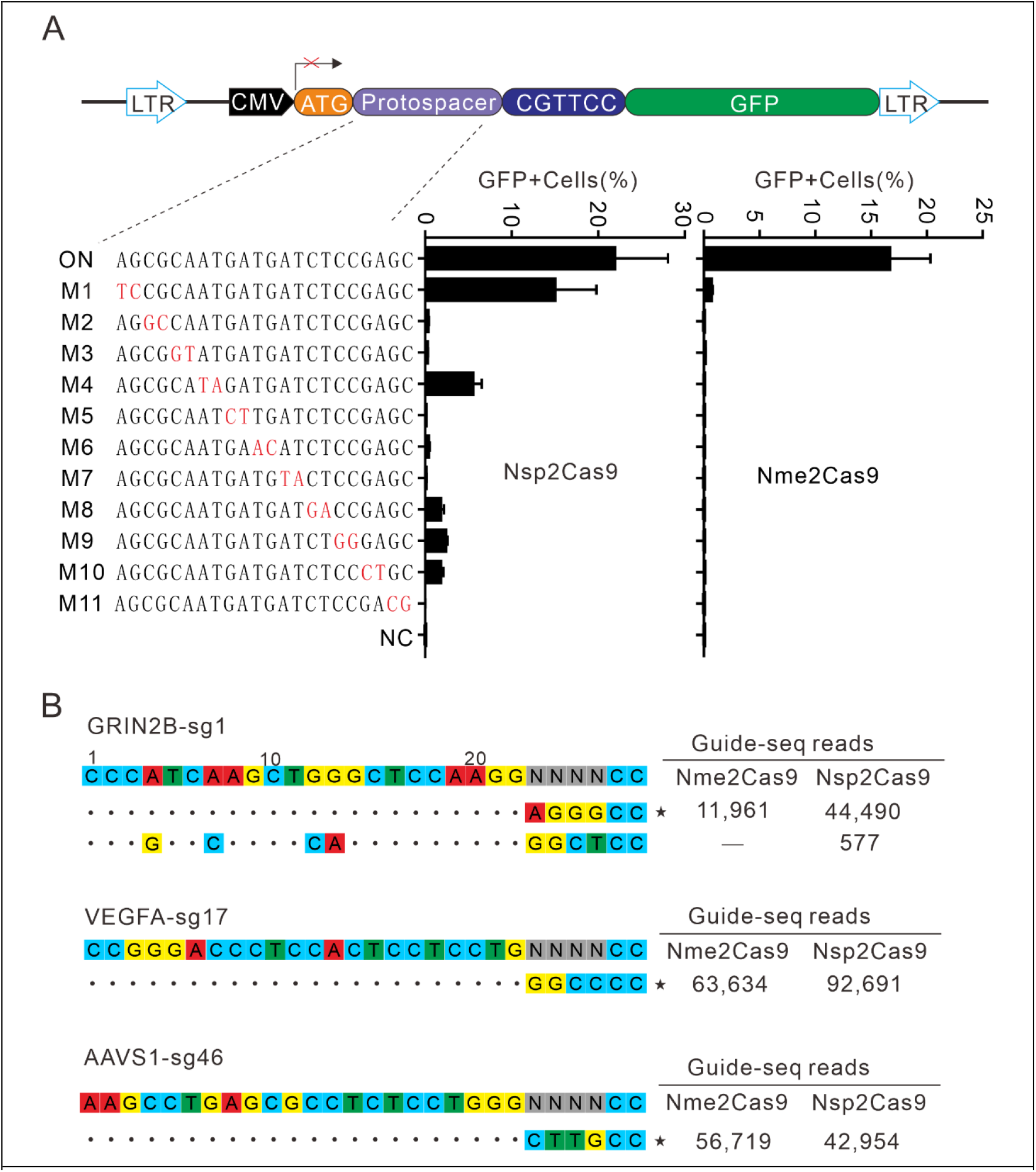
Analysis of Nsp2Cas9 specificity. **(A)** Analysis of Nsp2Cas9 and Nme2Cas9 specificity with a GFP-activation assay. A panel of sgRNAs with dinucleotide mutations (red) is shown below. The editing efficiencies reflected by ratio of GFP-positive cells are shown. Data represent mean ± SD for n=3 biologically independent experiments. **(B)** GUIDE-seq was performed to analyze the genome-wide off-target effects of Nsp2Cas9 and Nme2Cas9. On-target (indicated by stars) and off-target sequences are shown on the left. Read numbers are shown on the right. Mismatches compared to the on-target site are shown and highlighted in color.

To further compare the specificity of Nsp2Cas9 and Nme2Cas9, we performed genome-wide unbiased identification of double-stranded breaks enabled by sequencing (GUIDE-seq) (30). Three endogenous sites targeting GRIN2B, VEGFA and AAVS1 were selected. Five days after electroporation of the Cas9 expression plasmid with GUIDE-seq oligos into HEK293T cells, libraries were prepared for deep sequencing. The sequencing analysis revealed that both Cas9 nucleases could robustly generate indels at three on-targets, as reflected by the high GUIDE-seq read counts (Figure 5B). We only detected one off-target site for Nsp2Cas9 on the GRIN2B target (Figure 5B).

Simultaneously, we also analyzed Nsp2-SmuCas9 specificity by using the GFP-activation assay. Nsp2-SmuCas9 enzyme showed a minimal or background level of off-target effects with mismatched sgRNAs (Figure 6A). Next, GUIDE-seq was performed to test the specificity of Nsp2-SmuCas9. We selected three target sites containing N_4_C PAMs. The results revealed that one off-target cleavage occurred on the EMX1 target (Figure 6B). Altogether, these results demonstrated that the specificity of Nsp2Cas9 and Nsp2-SmuCas9 remained to be improved in future work.

**Figure 6.**
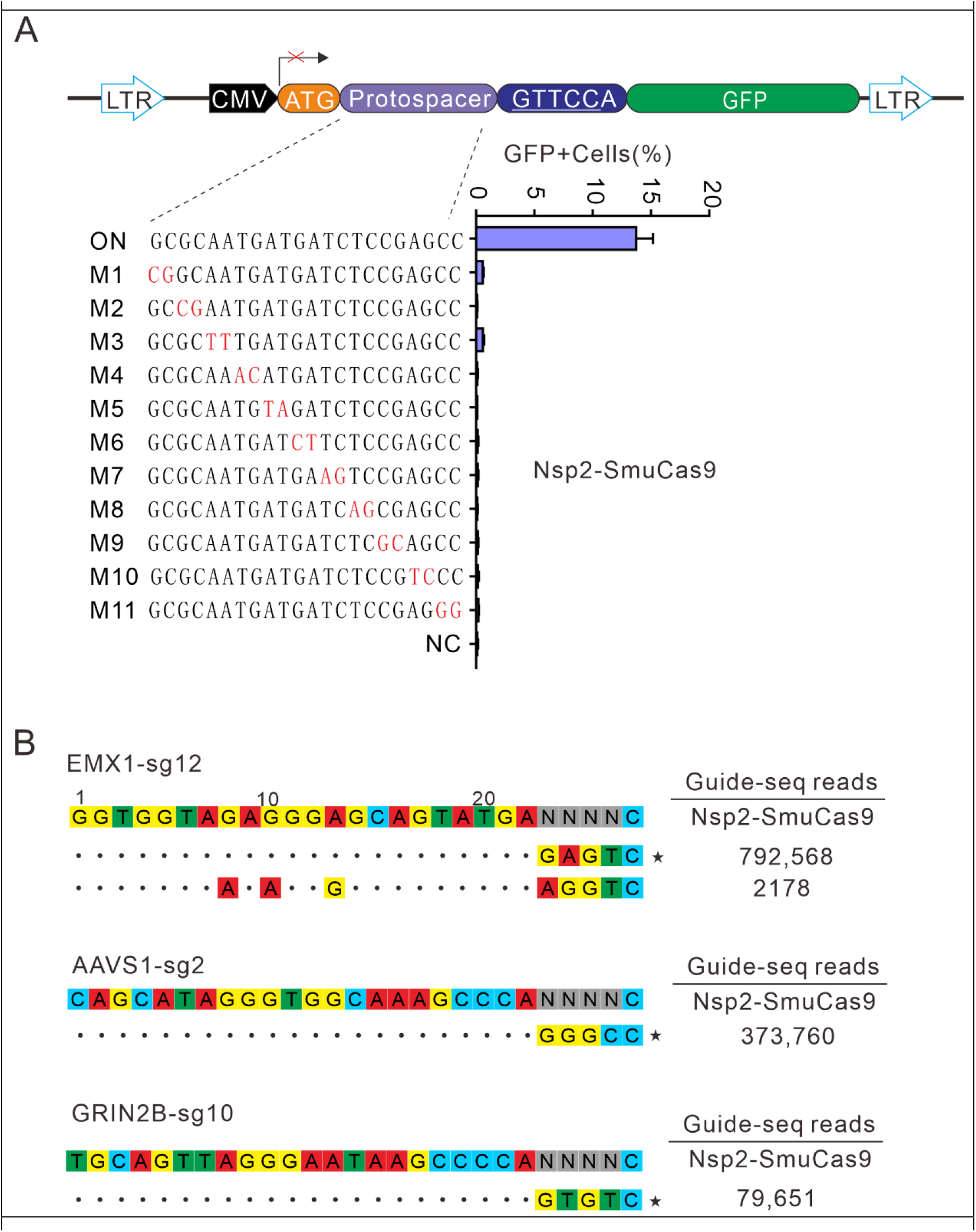
Analysis of Nsp2-SmuCas9 specificity. **(A)** Analysis of Nsp2-SmuCas9 specificity with a GFP-activation assay. A panel of sgRNAs with dinucleotide mutations (red) is shown below. The editing efficiencies reflected by ratio of GFP-positive cells are shown. Data represent mean ± SD for n=3 biologically independent experiments. **(B)** GUIDE-seq was performed to analyze the genome-wide off-target effects of Nsp2-SmuCas9. On-target (indicated by stars) and off-target sequences are shown on the left. Read numbers are shown on the right. Mismatches compared to the on-target site are shown and highlighted in color.

## Discussion

Type II CRISPR/Cas9, including subtypes II-A, -B and -C, is the most common class 2 system found in bacteria and archaea (31). We and others previously demonstrated that closely related type II-A orthologs preferred variable but purine-rich PAMs (17, 19-21, 32). The PAM diversity was also observed within the closely related type V-A orthologs, where Cas12a nucleases recognized the canonical TTTV (V=A/C/G) motif but notable deviations of nucleotide preference existed (33). In this study, we investigated the PAM diversity within closely related type II-C orthologs. We identified PAMs for 25 Nme1Cas9 orthologs, significantly extending the list of type II-C Cas9 PAMs. These PAMs included purine-rich (e.g., PaeCas9 and HfeCas9), pyrimidine-rich (e.g., Nsp2Cas9 and Nan1Cas9), and mixed purine and pyrimidine (e.g., HpiCas9 and NdeCas9) PAMs, which are more diverse than those identified from type II-A and type V-A orthologs. The PAM length also varied dramatically from a single nucleotide (e.g., NarCas9) to up to 11 nucleotides (e.g., CflCas9). Interestingly, phylogenetically conserved orthologs may recognize distinct PAMs. For example, PaeCas9 and AmiCas9 shared over 93% amino-acid identity. AmiCas9 recognized an N_4_RTTT PAM, while PaeCas9 recognized an N_4_GAAA PAM.

Our study enhanced ability to perform allele-specific genome editing. Allele-specific gene disruption through non-homologous end joining (NHEJ) is a potential strategy to treat autosomal dominant diseases (24, 34), where the causative gene is haplosufficient. Autosomal dominant diseases are mainly caused by single-nucleotide missense mutations (23). If missense mutations form novel PAMs, CRISPR/Cas9 nucleases can disrupt mutant alleles by a PAM-specific approach (24, 25). For clinical applications, it is crucial to develop a CRISPR toolbox capable of recognizing multiple PAMs. In this study, we identified 25 active Cas9 orthologs with different PAM requirement. The advantage of developing tools from closely related Cas9 orthologs is that they can exchange the PI domain. If an ortholog recognizes a particular PAM but does not work efficiently in human cells, we can use this ortholog PI to replace another ortholog PI to generate a chimeric Cas9. We used this strategy to generate an Nsp2-SmuCas9 that recognizes a simple N_4_C PAM. To our knowledge, the N_4_C PAM is the most relaxed PAM recognized by compact Cas9 orthologs identified to date. With further development, we anticipate that CRISPR tools from type II-C Cas9 orthologs can play a vital role in therapeutic applications.

## Materials and Methods

### Plasmid construction

#### Cas9 expression plasmid construction

The plasmid Nme2Cas9_AAV (Addgene #119924) was amplified by the primers Nme2Cas9-F/Nme2Cas9-R to obtain the Nme2Cas9_AAV backbone. The human codon-optimized Cas9 gene (Supplementary file 1) was synthesized by HuaGene (Shanghai, China) and cloned into the Nme2Cas9_AAV backbone by the NEBuilder assembly tool (NEB) according to the manufacturer’s instructions. Sequences of each Cas9 were confirmed by Sanger sequencing (GENEWIZ, Suzhou, China). The primer sequences for the Nme2Cas9_AAV backbone, variants of Nsp2Cas9 and chimeric Cas9 nucleases are listed in Supplementary file 1. Single-guide RNA sequences for each Cas9 are listed in Supplementary file 2.

### sgRNA expression plasmid construction

The sgRNA expression plasmids were constructed by ligating sgRNA into the Bsa1-digested U6-Nme2_scaffold plasmid, which is the same as Nme1-scaffold. The primer sequences and target sequences are listed in Supplementary file 3 and Supplementary file 4, respectively.

### Cell culture and transfection

The cell culture reagents were purchased from Gibco unless otherwise indicated. HEK293T, HeLa, SH-SY5Y, A375 and N2a cell lines were maintained in Dulbecco’s modified Eagle’s medium (DMEM) containing 10% fetal bovine serum (FBS) (Gibco) that was inactivated at 56 °C for 30 min and 1% penicillin-streptomycin (Gibco). All cell lines were cultured in a humidified incubator at 37 °C and 5% CO_2_.

For transient transfection, a total of 300 ng Cas9-expressing plasmid and 200 ng sgRNA plasmid were co-transfected into a 48-well plate. For Cas9 PAM sequence screening, 1.2×10^7^ HEK293T cells were transfected with 10 μg of Cas9 plasmid and 5 μg of sgRNA plasmid in 10-cm dishes.

### Flow cytometry analysis

Transfected library cells with a certain percentage of GFP-positive cells were collected by centrifugation at 1000 rpm for 5 min and resuspended in PBS. Then, GFP-positive cells were collected by flow cytometry and cultured in 6-well plates. Five days after culture, we extracted the genome and built deep sequencing libraries.

### PAM sequence analysis

Twenty-base-pair sequences (AAGCCTTGTTTGCCACCATG/GTGAGCAAGG GCGAGGAGCT) flanking the target sequence (GAGAGTAGAGGCGGCCACGACCTGNNNNNNNN) were used to fix the target sequences. CTG and GTGAGCAAGGGCG AGGAGCT were used to fix 8-bp random sequences. Target sequences with in-frame mutations were used for PAM analysis. The 8-bp random sequence was extracted and visualized by WebLogo (35) and a PAM wheel chart to identify PAMs (36).

### Genome editing and deep sequencing analysis of indels for endogenous sites

Cells were seeded into 48-well plates one day prior to transfection and transfected at 70–80% confluency using Lipofectamine 2000 (Life Technologies) following the manufacturer’s recommended protocol. For genome editing, 10^5^ cells were transfected with a total of 300 ng of Cas9 plasmid and 200 ng of sgRNA plasmid in 48-well plates. Five days after transfection, the cells were harvested, and genomic DNA was extracted in QuickExtract DNA Extraction Solution (Epicenter). To measure indel frequencies, the target sites were amplified by two rounds of nested PCR to add the Illumina adaptor sequence. The PCR products (∼400 bp in length) were gel-extracted by a QIAquick Gel Extraction Kit (QIAGEN) for deep sequencing.

### Western blot analysis

HEK293T cells were seeded into 24-well plates. The next day, the Cas9-expressing plasmid (800 ng) was transfected into cells using Lipofectamine 2000 (Invitrogen). Three days after transfection, cells were collected and resuspended in cell lysis buffer for Western blotting and IP (Beyotime) supplemented with 1 mM phenylmethanesulfonyl fluoride (PMSF) (Beyotime). Cell lysates were then centrifuged at 12,000 rpm for 20 min at 4 °C, and the supernatants were collected and mixed with 5x loading buffer followed by boiling at 95 °C for 10 min. Equal amounts of protein samples were subjected to SDS-PAGE, followed by transfer to polyvinylidene difluoride (PVDF) membranes (Merck, Darmstadt, Germany). The PVDF membranes were blocked with 5% BSA in TBST for 1 hour at room temperature and then incubated with the anti-HA antibody (1:1000; Abcam) and anti-GAPDH antibody (1:1000; Cell Signaling) at 4 °C overnight.

The membrane was washed three times in TBS-T for 5 min each time. The membranes were incubated in the secondary goat anti-rabbit antibody (1:10,000; Abcam) for 1 h at room temperature. The membranes were then washed with TBST buffer three times and imaged.

### Test of Cas9 specificity

To test the specificity of Nsp2Cas9 and Nsp2-SmuCas9, we generated a GFP reporter cell line with a fixed PAM (5’-CGTTCC-3’). HEK293T cells were seeded into 48-well plates and transfected with 300 ng of Cas9 plasmids and 200 ng of sgRNA plasmids by using Lipofectamine 2000. Five days after transfection, the GFP-positive cells were digested and centrifuged at 1000 rpm for 3 min, and the cells were resuspended in phosphate-buffered saline (PBS). Finally, the cells were analyzed on a Calibur instrument (BD). The data were analyzed using the FlowJo software.

### GUIDE-seq

We performed a GUIDE-seq experiment with some modifications to the original protocol, as described (37). On the day of the experiment, 2×10^5^ HEK293T cells per target site were harvested and washed in PBS and transfected with 500 ng of Cas9 plasmid, 500 ng of sgRNA plasmid and 100 pmol annealed GUIDE-seq oligonucleotides through the Neon™ Transfection System. The electroporation voltage, width, and number of pulses were 1,150 V, 20 ms, and 2 pulses, respectively. Genomic DNA was extracted with the DNeasy Blood and Tissue kit (QIAGEN) 6 days after electroporation according to the manufacturer’s protocol. The genome library was prepared and subjected to deep sequencing.

### Statistical analysis

All data are presented as the mean ± SD. Statistical analysis was conducted using GraphPad Prism 7. Student’s *t* test or one-way analysis of variance (ANOVA) was used to determine statistical significance between two or more than two groups, respectively. A value of *p* < 0.05 was considered to be statistically significant (\**p* < 0.05, \*\**p* < 0.01, \*\*\**p* < 0.001).

## Ethics approval and consent to participate

Not applicable.

## Consent for publication

Not applicable.

## Availability of data and materials

All data generated or analysed during this study are included in this published article and its supplementary information files.

## Author contributions

J.W., L.H., and S.G. performed experiments; J.L. and T.Q. analyzed the data; S.S., and Y.W. provide experimental guidance; Y.W. designed experiments and wrote the manuscript. All authors read and approved the final manuscript.

## Acknowledgements

Not applicable.

## Supplementary Information

**Figure 1-figure supplement 1.**
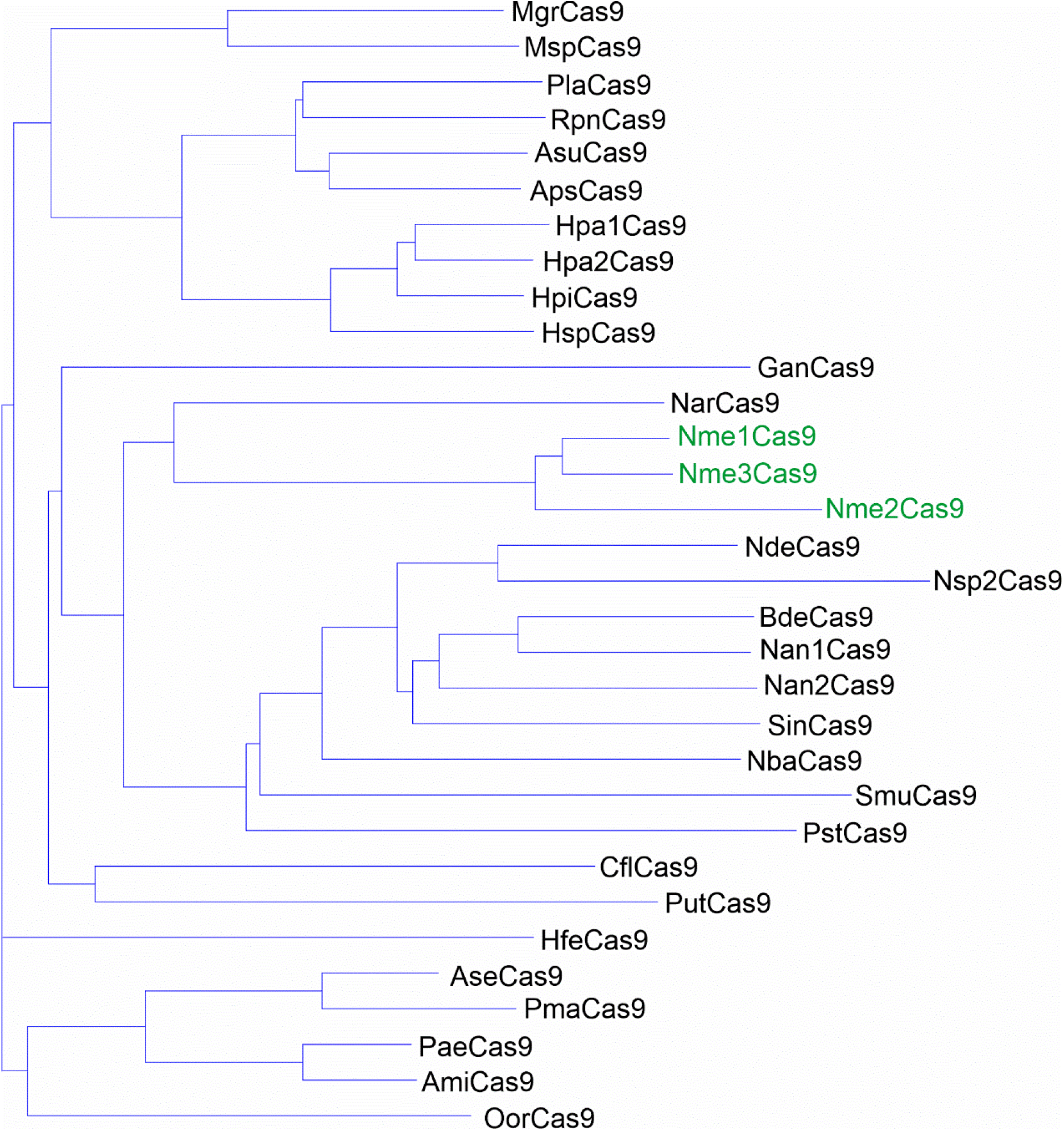
Phylogenetic tree of the selected Nme1Cas9 orthologs. The amino acid sequences of 29 selected Nme1Cas9 orthologs were aligned by Vector NTI. Nme1Cas9, Nme2Cas9, and Nme3Cas9 were used as reference and shown in green.

**Figure 1-figure supplement 2.**
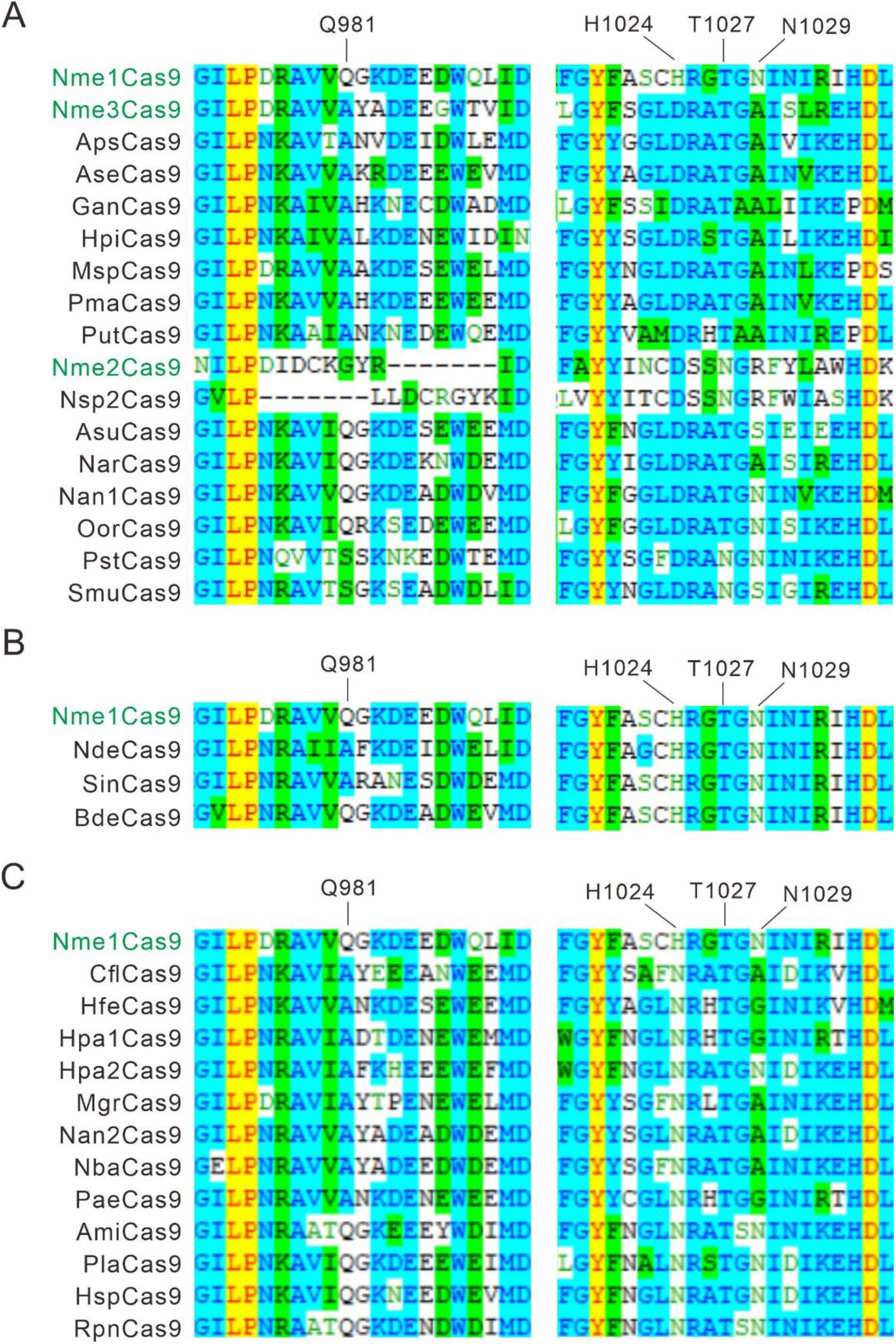
Alignment of the PI domain of Nme1Cas9 orthologs. The Nme1Cas9 orthologs contained aspartate **(A)**, histidine **(B)**, or asparagine **(C)** residues corresponding to the Nme1Cas9 H1024. The PI domains were aligned by Vector NTI. Amino acids crucial for PAM recognition are shown above. Nme1Cas9, Nme2Cas9, and Nme3Cas9 were used as reference and shown in green.

**Figure 1-figure supplement 3.**
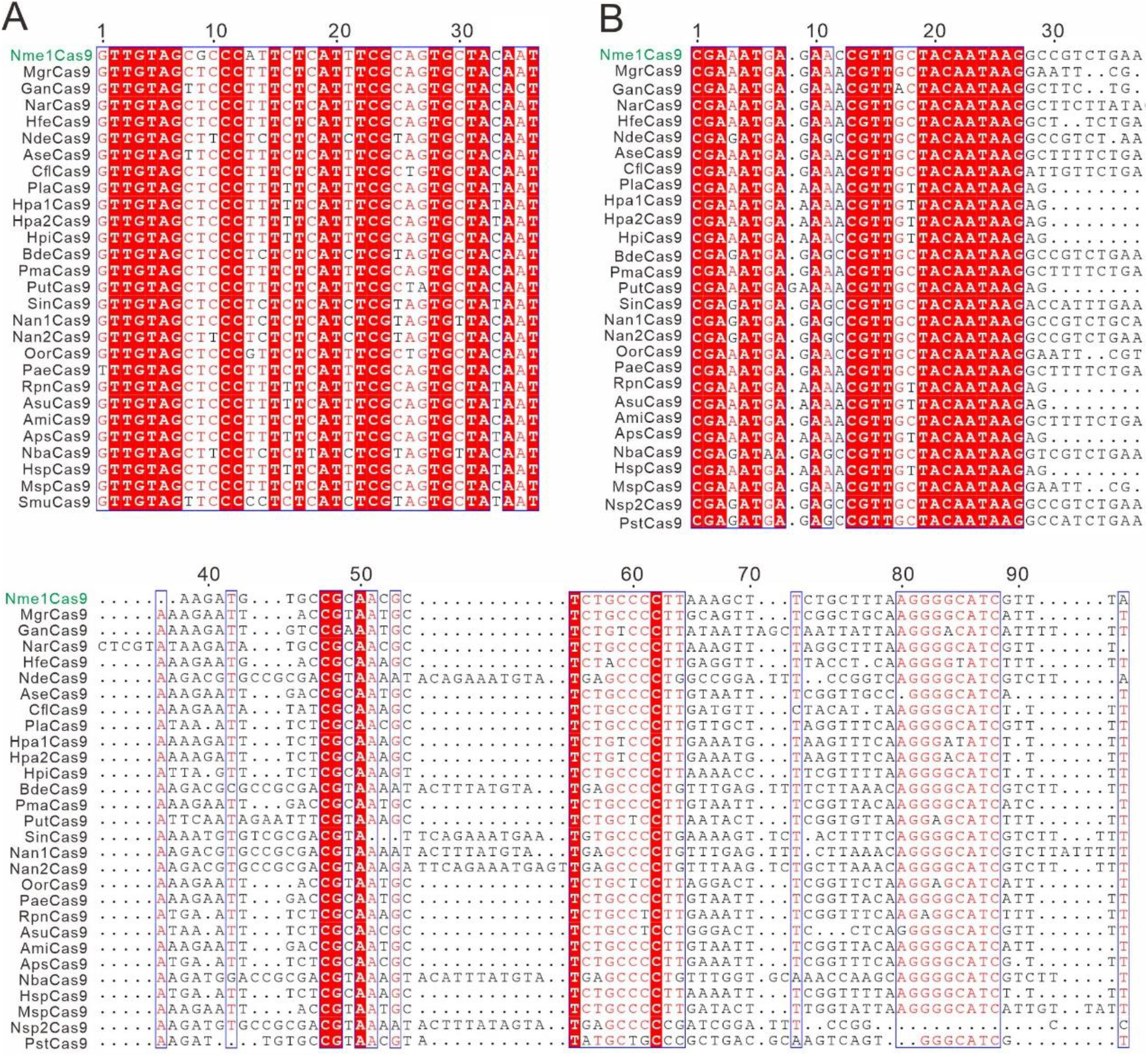
The alignment of direct repeats and tracrRNAs of Nme1Cas9 orthologs. **(A)** Alignment of direct repeat sequences for Nme1Cas9 orthologs is shown. **(B)** Alignment of tracrRNAs for Nme1Cas9 orthologs. Sequence alignment revealed that direct repeats and the 5’ end of tracrRNAs were conserved among Nme1Cas9 orthologs. Strict identical residues are highlighted with the red background and conserved mutations are highlighted with an outline and red font.

**Figure 1-figure supplement 4.**
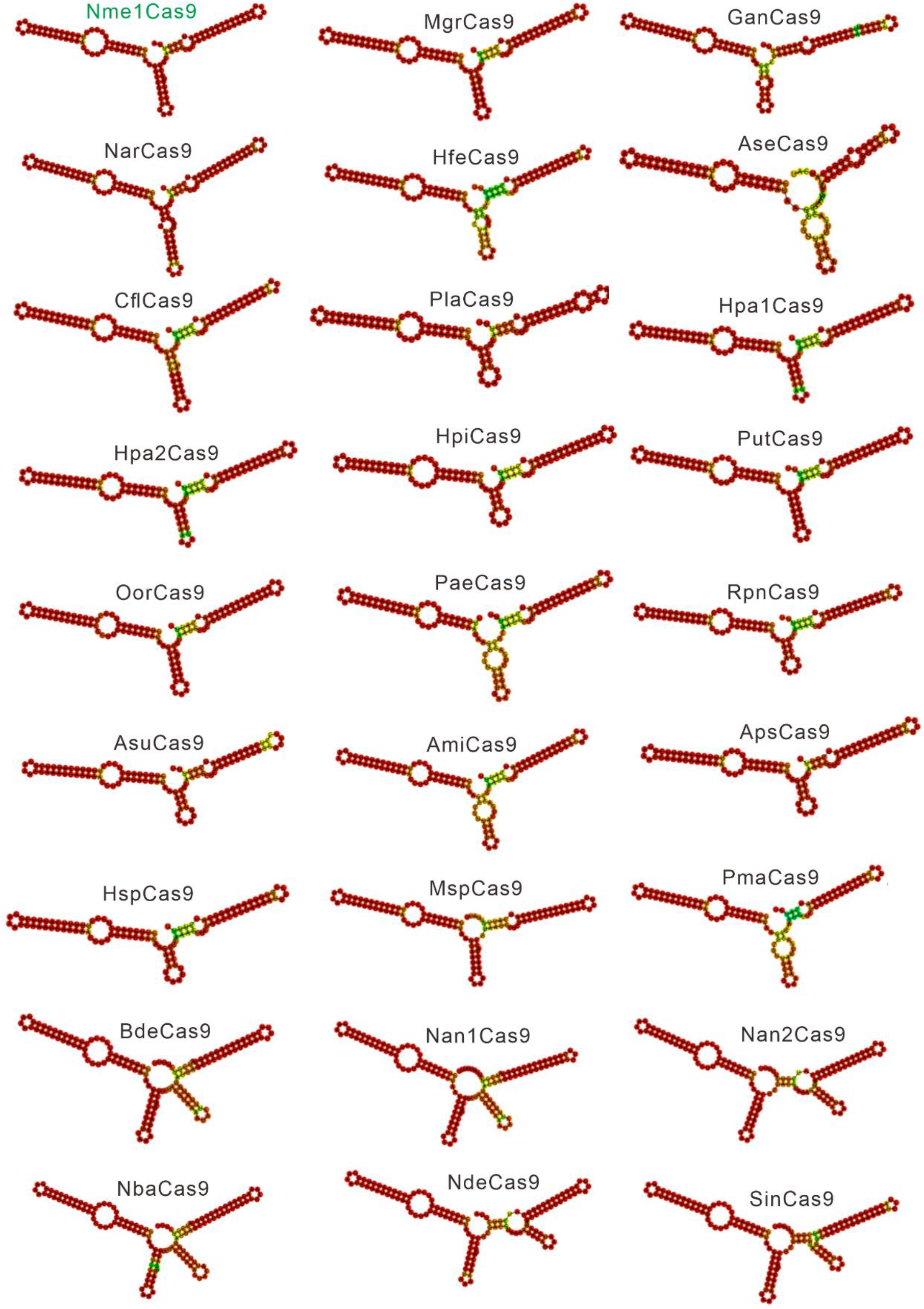
Single guide RNA (sgRNA) scaffolds of Nme1Cas9 orthologs. *In silico* co-folding of the crRNA direct repeat and putative tracrRNA shows stable secondary structure and complementarity between the two RNAs.

**Figure 2-figure supplement 1.**
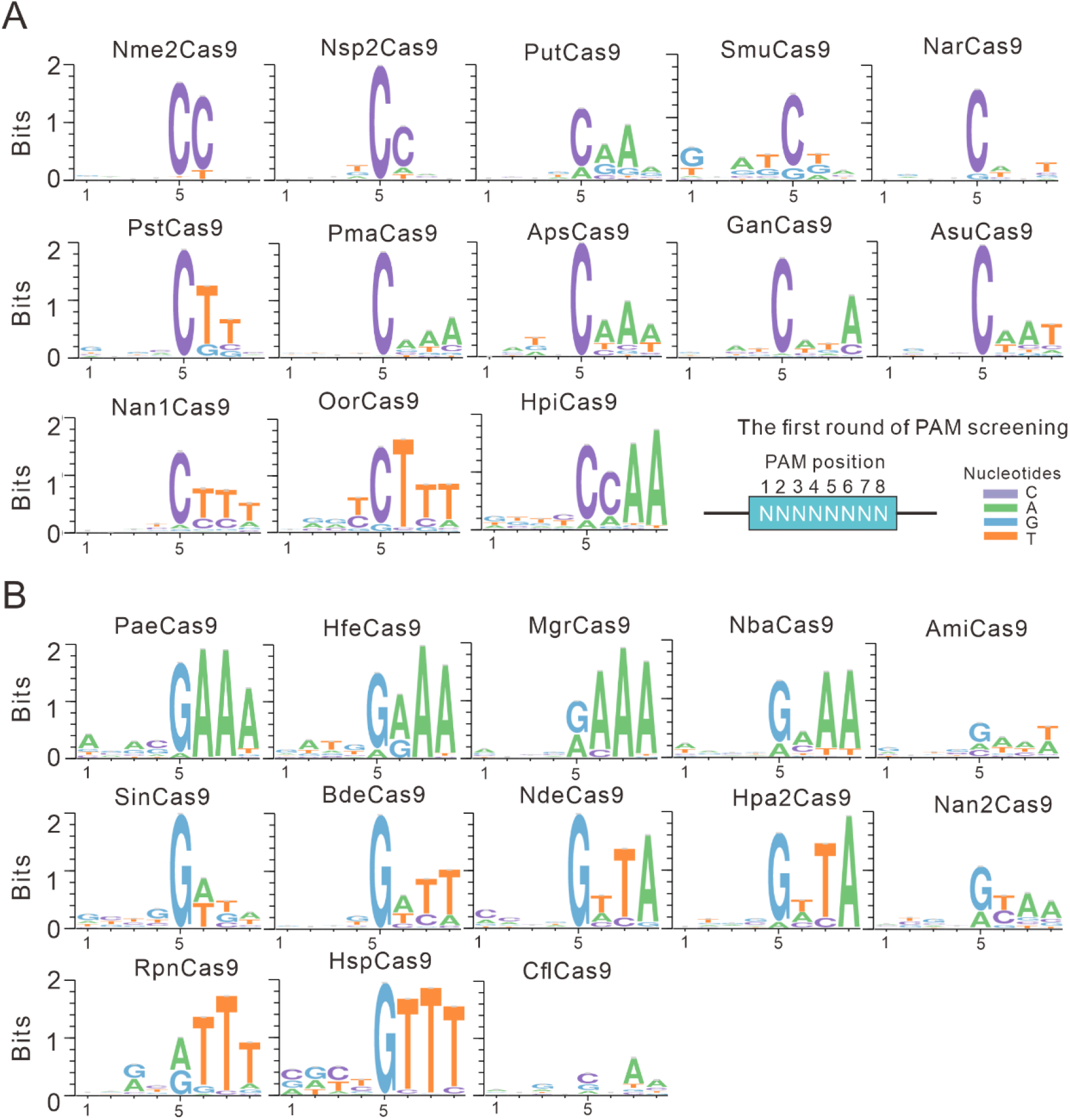
The WebLogo for each Cas9 nuclease. **(A)** PAMs for Nme1Cas9 orthologs containing an aspartate residue corresponding to the Nme1Cas9 H1024. PAM positions for the first round of screening are shown on the bottom right. **(B)** PAMs for Nme1Cas9 orthologs containing histidine, or asparagine residues corresponding to the Nme1Cas9 H1024.

**Figure 2-figure supplement 2.**
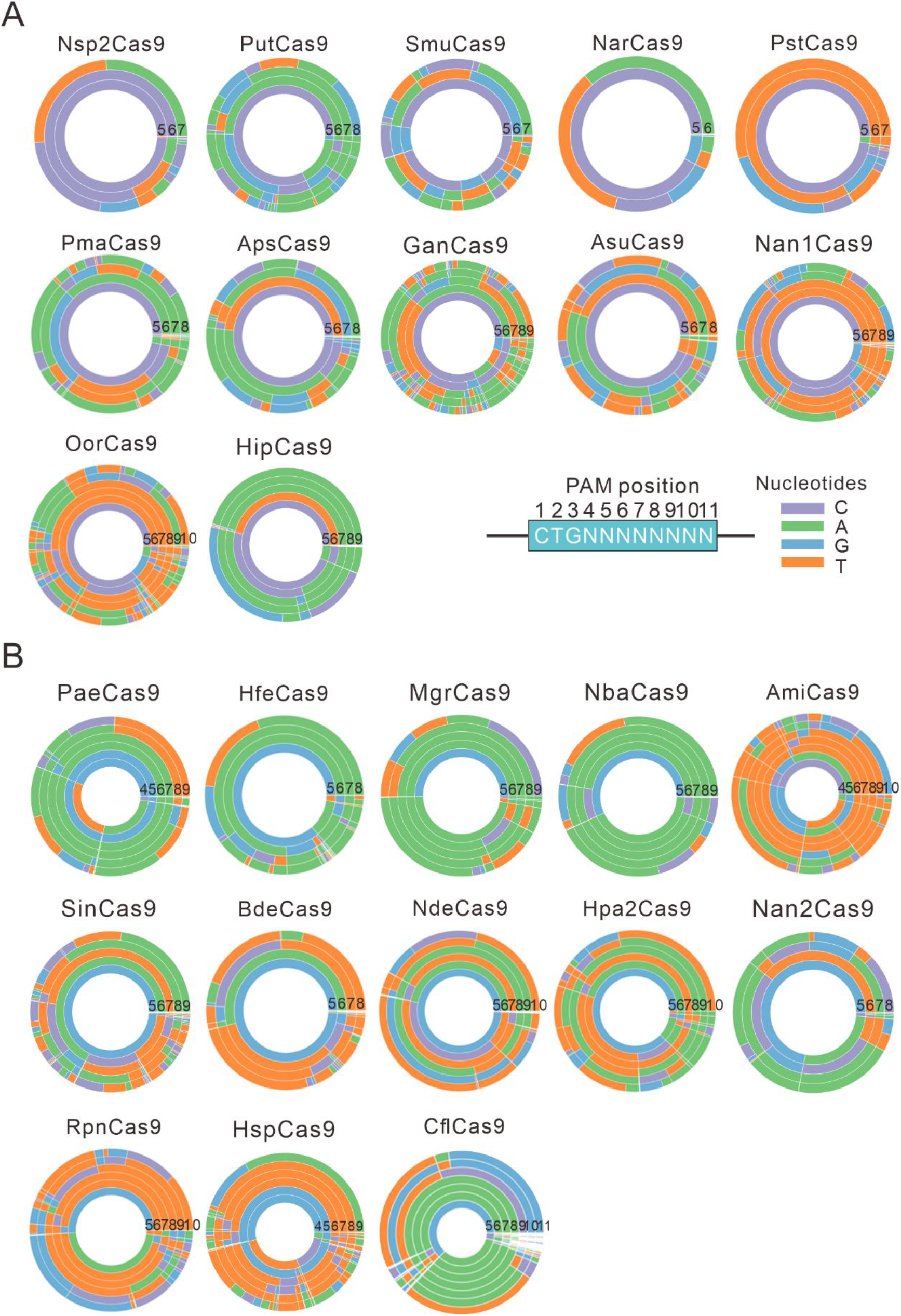
PAM wheels for Nme1Cas9 orthologs. **(A)** PAM wheels for Nme1Cas9 orthologs containing an aspartate residue corresponding to the Nme1Cas9 H1024. PAM positions in the screening assay are shown on the bottom right. (**B**) PAM wheels for Nme1Cas9 orthologs containing histidine, or asparagine residues corresponding to the Nme1Cas9 H1024. PAM wheels start in the middle of the wheel for the first 5’ base exhibiting sequence information.

**Figure 3-figure supplement 1.**
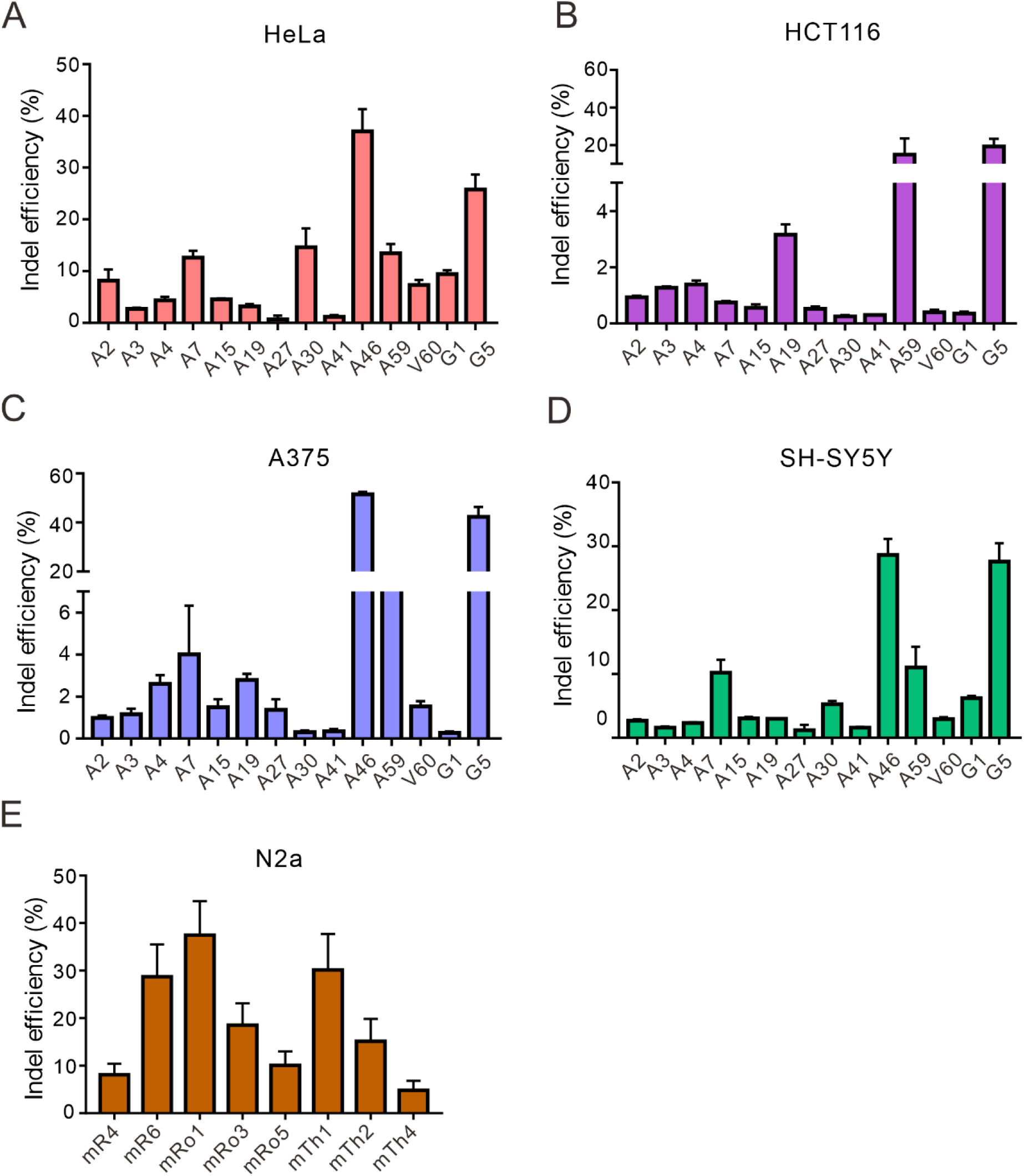
Nsp2Cas9 enables editing in different mammalian cells. Nsp2Cas9 enables editing in HeLa **(A)**, HCT116 **(B)**, A375 **(C)**, SH-SY5Y **(D)** and mouse N2a cells **(E)**. Data represent mean ± SD (n=3).

**Figure 3-figure supplement 2.**
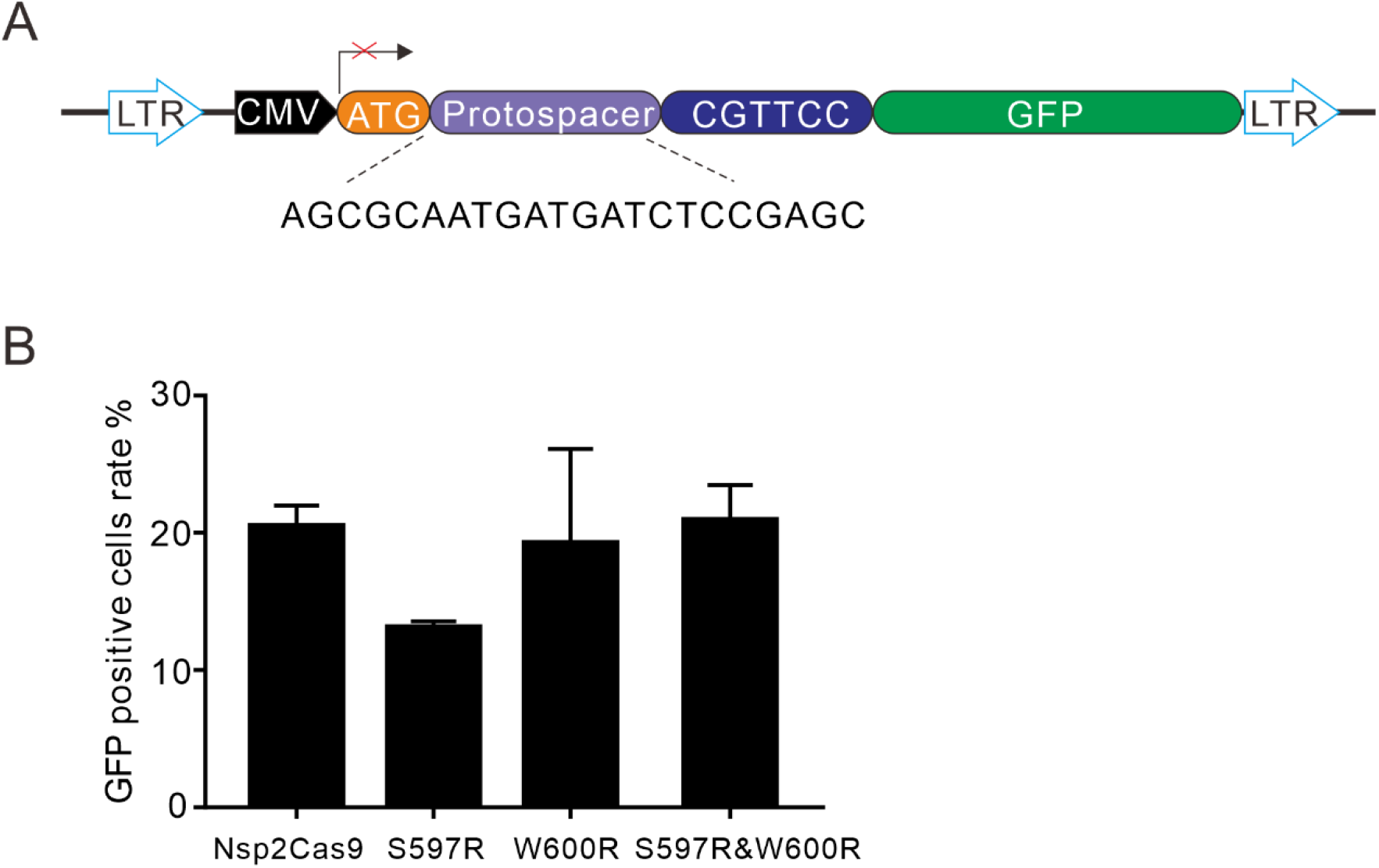
Rational engineering of Nsp2Cas9. **(A)** Schematic of the GFP-activation reporter construct for testing engineered Nsp2Cas9 activity. The protospacer sequence is shown below. **(B)** GFP-positive cells induced by the engineered Nsp2Cas9 variants. Data represent mean ± SD (n=3).

**Figure 3-figure supplement 3.**
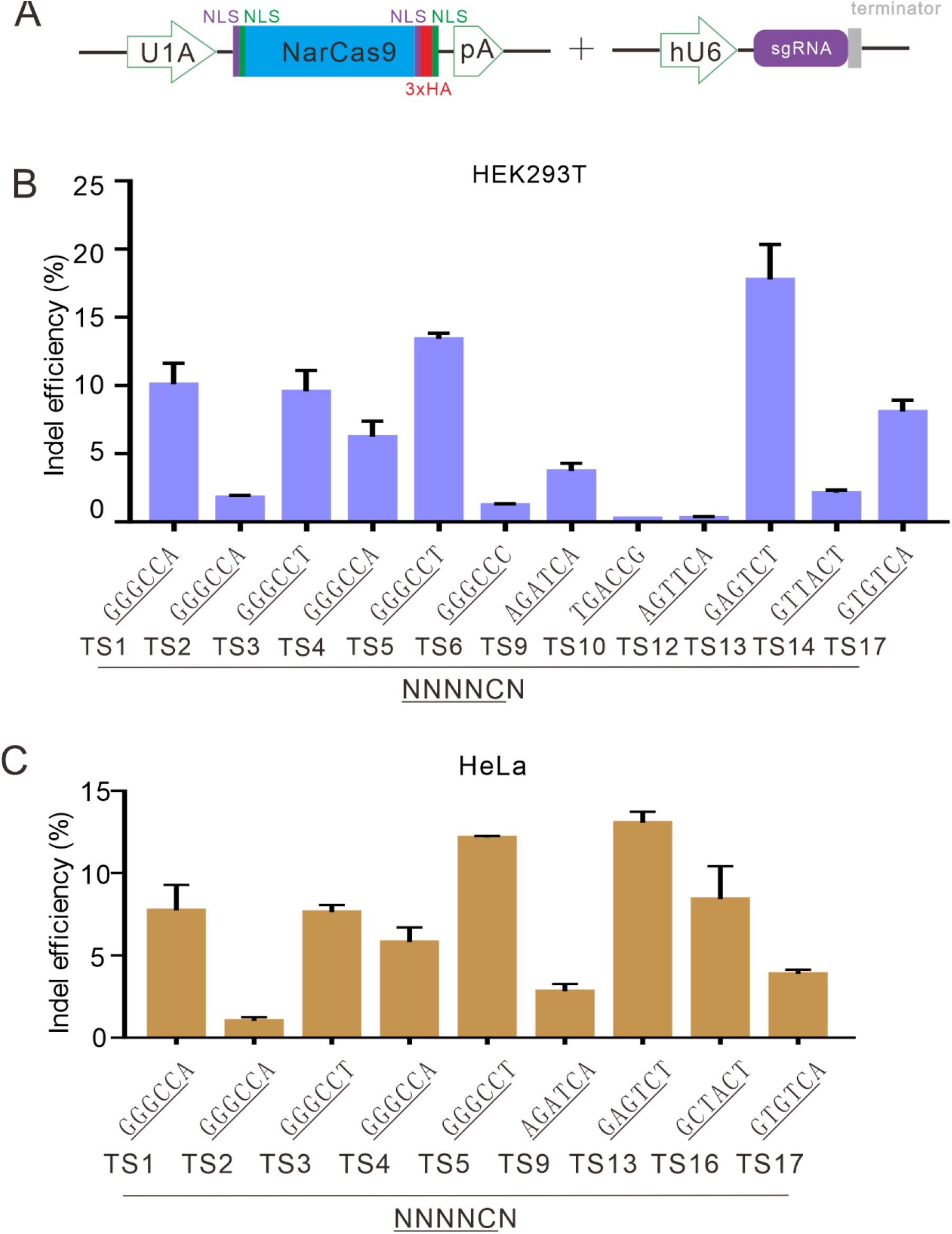
NarCas9 enables genome editing in mammalian cells. **(A)** Schematic of Cas9 and sgRNA expression constructs. U1A: U1A promoter; pA: polyA; NLS: nuclear localization signal; HA: HA tag. **(B)** NarCas9 enables genome editing in HEK293T cells. Data represent mean ± SD (n=3). **(C)** NarCas9 enables genome editing in HeLa cells. Data represent mean ± SD (n=3).

